# Div-Seq: A single nucleus RNA-Seq method reveals dynamics of rare adult newborn neurons in the CNS

**DOI:** 10.1101/045989

**Authors:** Naomi Habib, Yinqing Li, Matthias Heidenreich, Lukasz Swiech, John J. Trombetta, Feng Zhang, Aviv Regev

## Abstract

Transcriptomes of individual neurons provide rich information about cell types and dynamic states. However, it is difficult to capture rare dynamic processes, such as adult neurogenesis, because isolation from dense adult tissue is challenging, and markers for each phase are limited. Here, we developed Div-Seq, which combines Nuc-Seq, a scalable single nucleus RNA-Seq method, with EdU-mediated labeling of proliferating cells. We first show that Nuc-Seq can sensitively identify closely related cell types within the adult hippocampus. We apply Div-Seq to track transcriptional dynamics of newborn neurons in an adult neurogenic region in the hippocampus. Finally, we find rare adult newborn GABAergic neurons in the spinal cord, a non-canonical neurogenic region. Taken together, Nuc-Seq and Div-Seq open the way for unbiased analysis of any complex tissue.

---

Single cell RNA-Seq has greatly extended our understanding of heterogeneous tissues, including the CNS (Darmanis et al., 2015; Shin et al., 2015; Tasic et al., 2016; Thomsen et al., 2016; Usoskin et al., 2015; Zeisel et al., 2015), and is reshaping the concept of cell type and state. However, some key dynamic processes that occur in dense nervous tissues, such as adult neurogenesis, still remain challenging to study. First, single cell RNA-Seq requires enzymatic tissue dissociation, which damages the integrity of neurons (**Figure 1A**), compromises RNA integrity, and skews data towards easily dissociated cell types. This challenge is exacerbated as animals age, restricting this approach to fetal or young animals (Zeisel et al., 2015). Second, rare cells, such as adult newborn neurons found in the adult mouse hippocampus (Ming and Song, 2011), are difficult to capture because they require enrichment using specific tagging and sorting, for each phase of the dynamic neurogenesis process.

**Figure 1.**
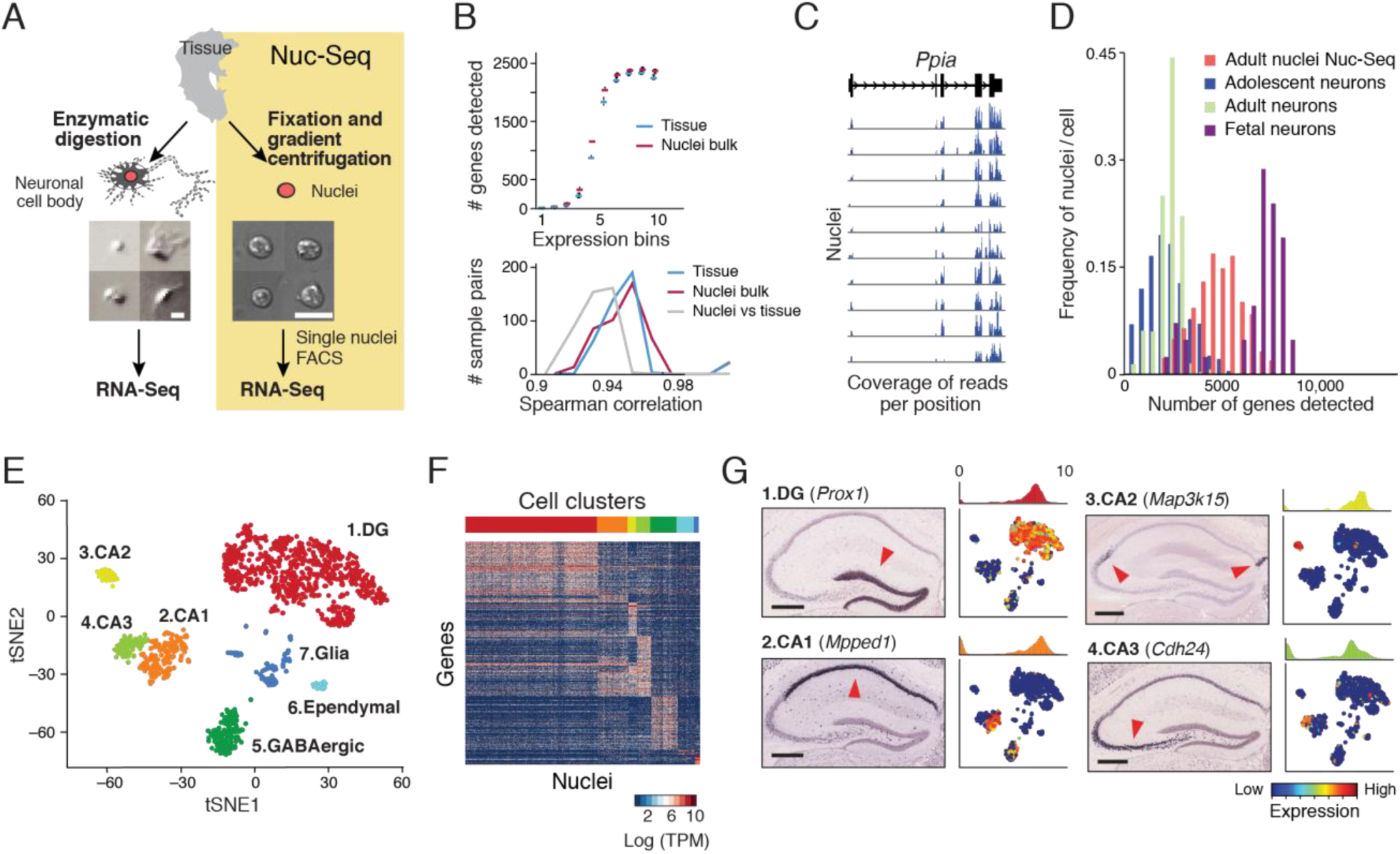
Single nuclei RNA-Seq (Nuc-Seq) identifies distinct cell types. (A) Single isolated nuclei (right) are more uniform than enzymatically dissociated single neuronal cell bodies (left) from adult mouse brain tissue. Shown are images of representative examples. Scale = 10p.m. Overview of the Nuc-Seq method (right): Dissected tissue is fixed, nuclei are isolated and sorted using FACS, and are then processed using the Smart-Seq2 RNA-Seq protocol(Picelli et al., 2013). (B) Nuc-Seq faithfully captures tissue RNA. Comparing Nuc-Seq on populations of nuclei and RNA-seq on tissue samples from the DG brain region. Shown are number of genes detected (TPM>3) per expression quantile (top) and distribution of pairwise spearman correlations across samples (bottom). (C) Nuc-Seq detects full-length, spliced transcripts in ten individual nuclei (rows). RNA-Seq read coverage at the *Ppia* genomic locus. (D) Nuc-Seq detects consistently higher number of genes (TPM/FPKM >3 or UMI>=1) compared to published single neuron RNA-seq in adolescent (Zeisel et al., 2015) or adult (Tasic et al., 2016) mice, but lower number than in fetal neurons (Thomsen et al., 2016). (E) Major cell types identified from Nuc-Seq data reflected by 7 major cell clusters. Shown is a 2-D non-linear embedding with 7 distinct clusters of 1,188 nuclei isolated from adult hippocampus. (F) Heatmap shows the expression of marker genes specific for each of the seven clusters across single nuclei (t-test FDR<0.05, log-ratio>1, across all pairwise comparisons). Top color bar matches cluster color in E. (G) Identification of DG granule cell, CA1, CA2, and CA3 pyramidal cell clusters. For each cluster, expression of marker genes is shown as: 1, ISH image in a coronal section of the hippocampus from (Lein et al., 2007) (arrowhead indicates high expression levels of marker gene); 2, histogram quantifying expression level across all nuclei in the relevant cluster; and 3, 2-D embedding of nuclei (as in E) showing relative expression level of the marker across all clusters. Scale = 400μm.

To address these challenges and study adult neurogenesis in an unbiased manner we set out to develop Div-Seq, a method to analyze single nuclei from recently dividing cells. Div-Seq relies on two advances. First, it uses **Nuc-Seq**, a high-throughput single-nucleus isolation and RNA-Seq method compatible with fresh, frozen, or fixed tissue. The uniform shape and fixation of the isolated nuclei (**Figure 1A**) combined with nuclei labeling (figure S1) enables enrichment of rare cell populations by fluorescent-activated cell sorting (FACS). Second, it uses unbiased labeling with 5-ethynyl-2’-deoxyuridine (EdU), which is incorporated into the DNA of dividing cells (Moore et al., 2015), and then uses Click-IT to fluorescently tag the isolated EdU labeled nuclei, which can be readily captured by FACS (figure S1).

Earlier studies have shown the feasibility of single neuronal nuclei RNA-seq (Grindberg et al., 2013; Krishnaswami et al., 2016; Swiech et al., 2015), however, it remains unclear whether the type and complexity of nuclear mRNA can be effectively used for sensitive classification of cell types and states in the CNS on large scale. Furthermore, given the relative low total amount and non-uniform distribution of RNA in neurons (nuclei, soma, axons, and dendrites), analysis of nuclei can introduce biases. We thus first tested nuclei RNA-Seq (Nuc-Seq) in bulk. Comparing RNA profiles of bulk tissue and populations of nuclei from the hippocampus dentate gyrus (DG) showed remarkable agreement, with similar RNA complexity and profiles (**Figure 1B**, in agreement with the previous observations (Grindberg et al., 2013)). Differential expression analysis shows that nuclear RNA enriches for long non-coding RNAs (figure S2). Thus, nuclear RNA contains as much information as tissue RNA, suggesting nuclear RNA-Seq does no introduce substantial biological biases.

Next, we analyzed 1,682 single nuclei from four hippocampal anatomical sub-regions (DG, CA1, CA2 and CA3) microdissected from adult mice, including genetically labeled and sorted GABAergic neurons nuclei that are of low abundance (~10% of total neuronal population (Hu et al., 2014), figure S1). Nuc-Seq detected 5,100 expressed genes per nucleus on average (**Figure 1C-D**), with comparable quality metrics to single-cell (non-neuron) RNA-Seq libraries (figure S2) and better library complexity (1.9-fold on average) compared to published single neuron RNA-Seq data (Shin et al., 2015; Tasic et al., 2016; Zeisel et al., 2015), across a wide range of expression levels (**Figure 1D** and figure S3). The range of transcripts detected was significantly improved compared to that of previously analyzed single nuclei (Grindberg et al., 2013) (two nuclei, figure S3). Finally, the complexity of Nuc-Seq libraries were similar in young (1 month), adult (3 months), and old (2 years) mice (figure S2), demonstrating robustness across animal ages. Thus, Nuc-Seq generated high quality data, exceeding the sensitivity of current single neuron RNA-seq.

Nuc-Seq analysis sensitively identified both major cell types and refined sub-types. Cluster analysis of Nuc-Seq data revealed seven major clusters of cells with distinct gene expression patterns (**Figure 1E-G**, figure S4 and S5, and table S1) that clearly correspond to known cell types and major anatomical distinctions in the hippocampus. Cluster identities were consistent with our microdissection scheme, and their gene expression patterns globally agreed with Allen Brain Atlas ISH data (Allen ISH (Lein et al., 2007), figure S5). Iterative re-clustering of the glia nuclei (cluster 7 in **Figure 1E** and figure S6) recovered five known glial cell sub-types (Zhang et al., 2014b), and averaged expressions across each sub-cluster well-correlated with published population RNA-Seq data (Zhang et al., 2014b) (figure S6 and table S1).

We captured finer distinctions between closely related cell types using a new clustering algorithm, biSNE (biclustering on Stochastic Neighbor Embedding) (**Figure 2** and figure S7-8), which improved upon current methods (Anders and Huber, 2010) (figure S7). The biSNE analysis partitioned the GABAergic neurons into eight sub-clusters (**Figure 3A**), each with unique expression of individual or pairs of canonical interneuron marker genes, such as *Pvalb* and *Htr3a* (**Figure 3B**). We validated the expression patterns of GABAergic markers by double fluorescent RNA *in situ* hybridization (dFISH) (**Figure 3C** and figure S9). We further characterized the sub-clusters by differential gene expression analysis (table S2), revealing for example that the calcium channel *Cacna1i* is specifically expressed in *Pvalb* or *Sst* positive GABAergic neurons (figure S8 and table S2).

**Figure 2.**
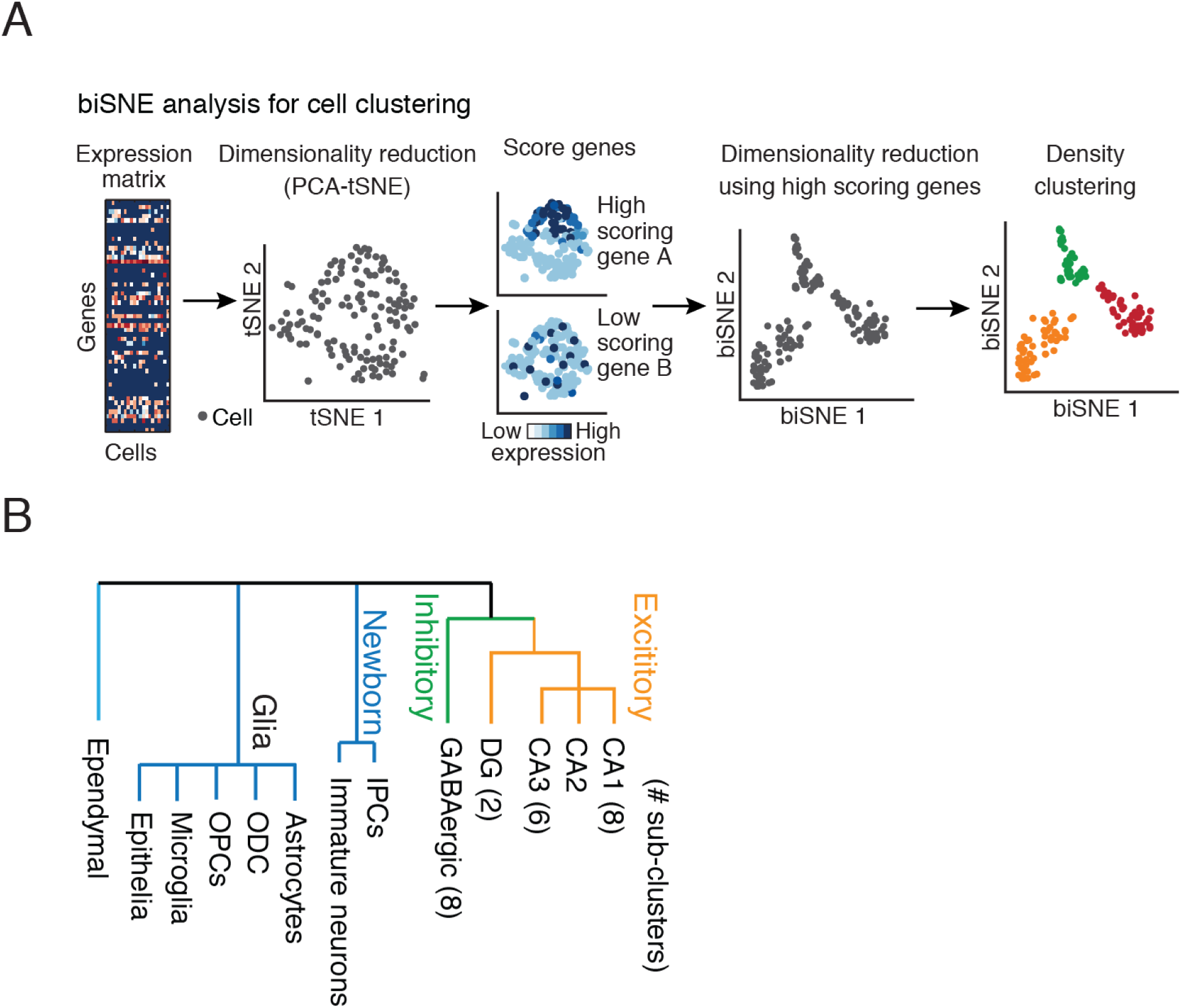
biSNE algorithm captures finer distinctions between cell types. (A) BiSNE algorithm. Top row, left to right: BiSNE takes as input an expression matrix of genes (rows) across nuclei (or cells, columns). It generates a 2-D plot of nuclei by dimensionality reduction using PCA followed by tSNE non-linear embedding, and then scores each gene by their expression across the 2-D plot, such that genes expressed in nuclei in proximity on the 2-D plot (dark blue points, top) are high scoring, whereas those expressed in nuclei scattered across the plot (dark blue points, bottom) are low scoring. Next, it takes an expression matrix of only high scoring genes (heatmap, genes (rows) across all nuclei (columns)), and repeats the dimensionality reduction. BiSNE is followed by density clustering (colored). (B) BiSNE sub-clustering. Dendrogram of all nuclei clusters along with the number of sub-clusters found by biSNE. NPC: neuronal precursor cells, ODC: oligodendrocytes, ASC: astroglia, OPC: oligodendrocyte precursor cell.

**Figure 3.**
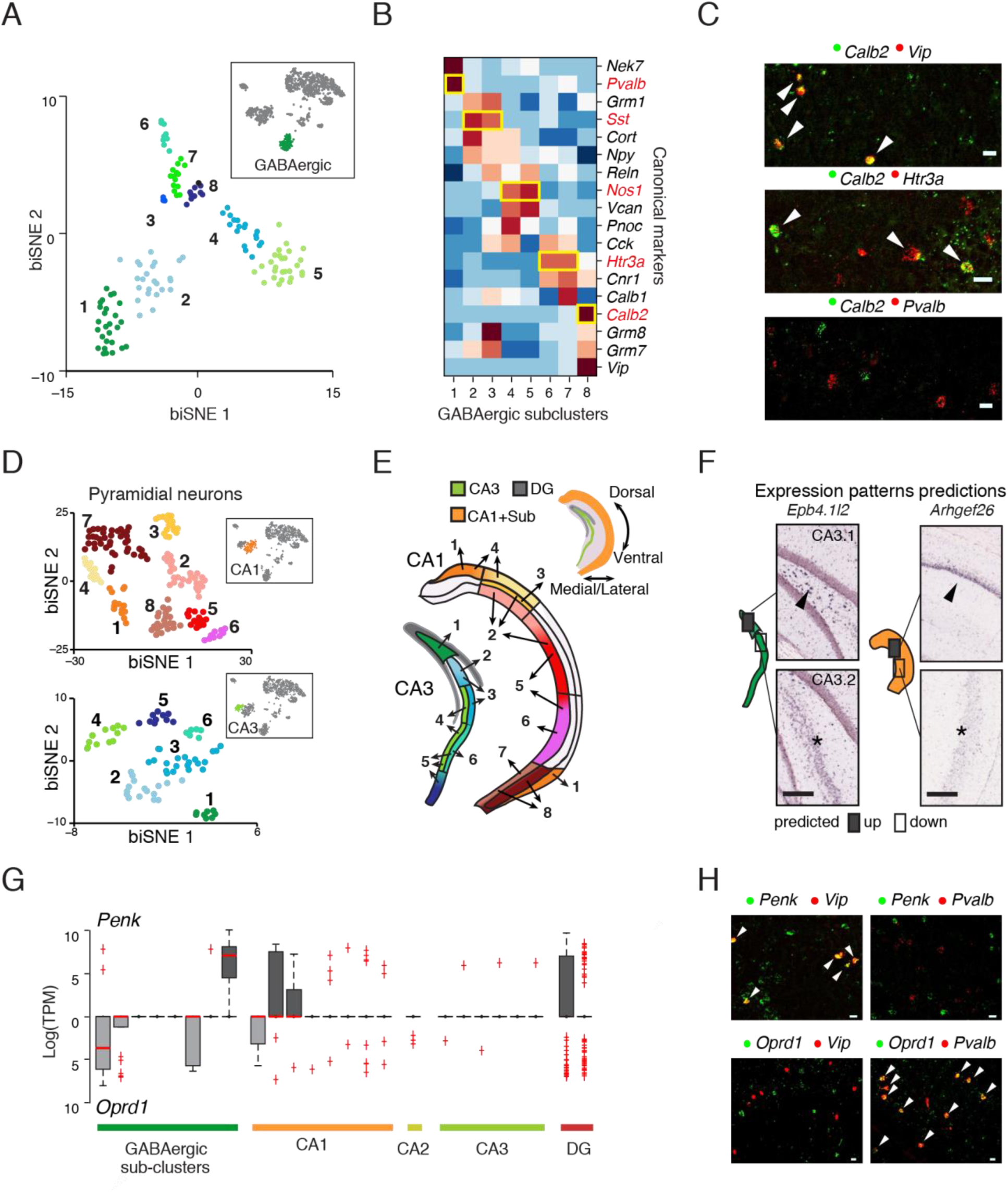
Nuc-Seq and biSNE distinguish cell subtypes and transcription patterns. (A) Subclusters of GABAergic interneurons identified by biSNE. Shown is a biSNE 2-D embedding of GABAergic nuclei with 8 sub-clusters. Top insert: the GABAergic cluster within all other nuclei from **Figure 1E**. (B) Sub-clusters are characterized by a combination of canonical marker genes. Heat map with averaged expression of canonical neuron markers (rows) across GABAergic subclusters (columns). (C) Double fluorescent RNA *in situ* hybridization (dFISH) of marker genes validating the expression pattern shown in (B). Co-expression of genes indicated by arrowheads. Scale = 20μm. (D) Pyramidal CA1 and CA3 biSNE sub-clusters. Shown is a biSNE 2-D embedding of the CA1 (top) and CA3 (bottom) pyramidal nuclei with 8 and 6 sub-clusters, respectively. Top insert: the CA1 cluster (orange) within all other nuclei from **Figure 1E**. (E) Spatially resolved pyramidal neuron populations in CA1 and CA3. Top: Schematics of hippocampus coronal section with CA1 (including subiculum), CA3 (including the hilus) and DG. Bottom: Registration of CA1 (right) and CA3 (left) pyramidal sub-clusters to subregions, using a map of landmark gene expression patterns from ISH data. Sub-cluster assignments are numbered and color code as in (D). Scale = 200μm. (F) Example of validation of spatial assignments of CA1 and CA3 pyramidal sub-clusters. Predictions (left illustrations; boxes showing predicted differential expression regions) match with Allen ISH data (Lein et al., 2007) (right; arrowhead: high expression; asterisk: low expression) in pairwise comparison of genes differentially expressed between two sub-clusters. (G) Distribution of expression of *Penk* (facing up) and *Oprd1* (facing down) across each neuronal sub-cluster. Box plots show the median (red), 75% and 25% quantile (box), error bars(dashed lines), and outliers (red cross). (H) dFISH of GABAergic cluster marker genes (*Vip* and *Pvalb*) with *Penk* or *Oprd1*, validating their mutual exclusive expression across GABAergic sub-clusters. Co-expression of genes indicated by arrowheads. Scale = 20μm.

Nuc-Seq also distinguished between spatial sub-regions with divergent transcriptional profiles. biSNE analysis of NucSeq data partitioned glutamatergic cells from CA1, CA3, and DG into 8, 6, and 2 sub-clusters, respectively (**Figure 3D** and figure S10). Analysis of sub-cluster specific gene expression highlighted several known landmark genes that exhibit spatially restricted expression patters in sub-regions of the hippocampus, indicating a correspondence between hippocampal sub-regions and sub-clusters of glutamatergic nuclei. We then used the spatial expression patterns (Lein et al., 2007) of these landmark genes to map sub-clusters in CA1, CA3, and DG to distinct spatial sub-regions (**Figure 3E** and figure S11, S12, S13). Notably, multiple sub-regions were assigned different, yet partially overlapping, sets of sub-clusters, indicating a gradual transition of transcriptional profiles between neighboring hippocampal sub-regions (**Figure 3E**). Other sub-regions were assigned to a single sub-cluster; in particular, a rare set (7%) of sparse neurons in the dorsal lateral outskirts of the CA1 (**Figure 3E**). To validate our mapping, we selected genes that were not used in the spatial mapping, and confirmed their predicted expression patterns in sub-regions of the hippocampus using the Allen ISH dataset (**Figure 3F** and figure S14). Previous studies using single-neuron RNA-Seq in CA1 reported two cell clusters that do not match spatial position (Zeisel et al., 2015) (figure S15), whereas our spatial mapping of Nuc-Seq data corresponds to continual transcriptional transitions within CA1 and CA3 regions, adding to the growing evidence (Cembrowski et al., 2016; Strange et al., 2014) that cellular diversity is not always partitioned into discrete sub-types.

We identified marker genes that are specifically associated with cell type and/or position (tables S1-S3). For example, *Penk*, which encodes an opioid neuropeptide (Enkephalin), and its receptor *Oprd1* (Roques et al., 2012), were selectively expressed in mutually exclusive sub-clusters of cells (**Figure 3G**): *Penk* was expressed in pyramidal neurons from dorsal CA1 and in *Calb2+/Vip+* GABAergic neurons, while *Oprd1* was expressed in subicular glutamatergic neurons and in *Pvalb+* and *Htr3a+/Cck+* GABAergic neurons. We validated the mutually exclusive expression pattern of *Penk* and *Oprd1* in GABAergic neurons by dFISH and their spatial expression pattern within the hippocampus by ISH (**Figure 3H** and figure S16-17). In DG granule neurons, we found mutually exclusive expression of *Penk* in a small subset of cells (162/674) and of *Cck* neuropeptide (Cholecystokinin) in all others (figure S16 and supplementary note), which we validated by quantitative PCR (figure S16). Previous work showed that Enkephalin is secreted to the extracellular space (Roques et al., 2012), and its signaling may not require synaptic connection. Thus, the cell-type specific expression of *Penk* and *Oprd1* points to putative cell types and spatial positions involved in the Enkephalin signaling within the hippocampal circuitry.

We next combined Nuc-Seq with EdU labeling of dividing cells, in a method we call Div-Seq (**Figure 4A**). In contrast to commonly used genetic labeling techniques (Llorens-Bobadilla et al., 2015; Shin et al., 2015; Zhao et al., 2010), which might be limited to specific cell types and requires cell types or developmental stage marker genes (Llorens-Bobadilla et al., 2015; Shin et al., 2015; Zhao et al., 2010), EdU tags newly synthesized DNA in dividing cells at a given time window, allowing for unbiased isolation of nuclei of neural stem cells and their progeny with high temporal resolution. To study transcriptional dynamics during adult neurogenesis in the DG, one of the canonical neurogenic sites in the mammalian CNS (Ming and Song, 2011), we used Div-Seq to isolate nuclei at 2 and 14 days after cell division, representing neural precursor cells (NPC), neuroblasts, and immature neuronal stages of adult neurogenesis, respectively (Ming and Song, 2011) (**Figure 4B** and figure S18). Div-Seq enriched for a broad range of newborn cells (figure S16). Expression of stage-specific marker genes confirmed that Div-Seq captured cells at distinct stages: 2-day labeled nuclei expressed NPC (*Tbr2/Eomes*) and neuroblast (*Sox4*) markers, whereas 7-day and 14-day nuclei expressed immature neuronal markers (*Sox11* and *Dcx*) (**Figure 4C**). Of note, *Dcx* a commonly used marker gene for immature neurons was expressed in all mature GABAergic neurons in the hippocampus (figure S18), highlighting the limits of using single marker genes to identify cell types.

**Figure 4.**
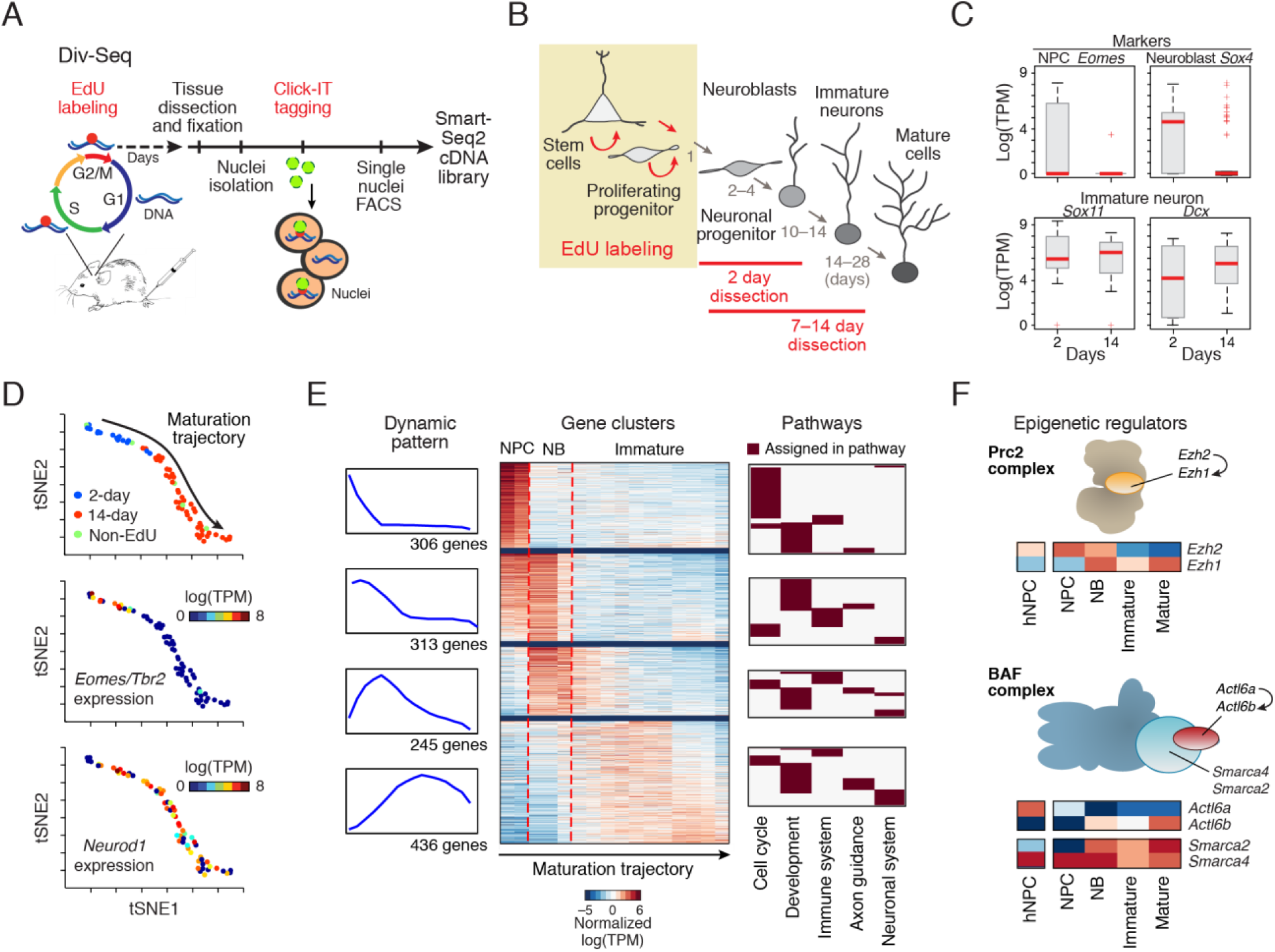
Transcriptional dynamics of adult neurogenesis revealed by Div-Seq. (A) Schematics of Div-Seq method. EdU is injected to mice and incorporates into the DNA of dividing cells (Moore et al., 2015). After isolation, EdU labeled nuclei are fluorescently tagged and captured by FACS for single nuclei RNA-Seq. (B) Schematics of adult neurogenesis in the dentate gyrus(Ming and Song, 2011). Timing of EdU labeling (tan box) and nuclei isolation are marked. (C) Div-Seq captured cells expressing known markers of neuronal precursors, neuroblasts and immature neurons. Box plots for the 2 days (2d) and 14 days (14d) EdU labeled nuclei (excluding nuclei classified as non-neuronal). Boxplots shown as in **Figure 3G**. (D) Newborn cells form a continuous trajectory. All panels show 2-D embedding of 2d labeled nuclei, 14d labeled nuclei and nuclei from unbiased survey. Nuclei are colored by source (top), by *Eomes*/*Tbr2* marker gene expression (middle), or by *Neurod1* marker expression (bottom). Trajectory directionality is chosen by the position of the 2d labeled neurons and known marker genes. (E) Dynamic gene expression clusters. Four clusters are shown from top to bottom. Left: Running average expression level of the genes in each cluster over the nuclei ordered along the trajectory (as in D). Middle: a heatmap of running average expression of all genes along the trajectory. Red lines mark the transcriptional switches from neuronal precursor cell (NPC) to neuroblast (NB), and from NB to immature neuron. Right: proportions of genes assigned to five major biological pathways (F) Changes in the composition of the Polycomb Complex (Prc2, top) and the BAF (SWI/SNF) complex (bottom). For each complex, schematics of the complex is shown, and the heatmap of average expression of genes in NPC(NP), NB, immature neurons and mature granule DG cells, and compared to human NPCs (hNPC, absolute log (TPM)).

Clustering analysis of neuronal lineage nuclei placed the newborn neurons on a continuous trajectory. The order of nuclei along the trajectory matched the EdU labeling time, from 2-day to 14-day labeled nuclei, with partial overlap, and a few nuclei from our unbiased survey of nuclei spread throughout (**Figure 4D**). Expression patterns of known neurogenesis genes along the trajectory recapitulated their known dynamics (Schouten et al., 2012; Shin et al., 2015; Tasic et al., 2016) and correctly captured the measured expression of nuclei at an intermediate time point of 7 day post EdU injection (figure S18), indicating that the trajectory indeed captured the maturation process.

To further characterize the transcriptional transitions of newborn neurons, we used biSNE to identify genes with dynamic expression patterns along the neurogenesis trajectory (**Figure 4E** and table S4), clustered genes by their expression patterns, and tested for enriched genetic pathways in each cluster. We found two major coordinated transcriptional switches, involving hundreds of genes and aligning with the known transitions from NPC, through neuroblasts, to immature neurons: (i) from proliferation (cell-cycle exit) to neuronal differentiation (consistent with previous reports (Shin et al., 2015)), and (ii) from differentiation to neuronal maturation (**Figure 4E**).

We identified transcription factors (TFs) and chromatin regulators whose expression is coordinated with these two transcriptional switches (figure S19). For the Polycomb Complex (Prc2), we observed an expression switch between *Ezh2* (expressed in NPCs consistent with previous reports (Zhang et al., 2014a)) and its paralog *Ezh1* (**Figure 4F**); for the BAF (mammalian SWI/SNF) complex, we observed an expression switch of *Actl6a/Baf53a* to its paralog *Actl6b/Baf53b* (Wu et al., 2007) and a late induction of BAF components (*e.g. Smarca2/BAF190b*, **Figure 4F** and figure S19). These expression patterns are consistent with single cell RNA-Seq of mouse NPCs (Shin et al., 2015) and human NPCs (Andrew Venteicher) (**Figure 4F** and figure S19).

Div-Seq provides a unique opportunity to profile the transcriptional program underlying neuronal maturation. We found differentially expressed genes between immature and mature DG granule neurons (t-test FDR q-value<0.01; figure S20, and differentially expressed splice isoforms, table S5), enriched for expected molecular pathways (q-value<0.01, figure S20), such as semaphorin signaling (Miller et al., 2013) (figure S20) and lipid metabolism (Knobloch et al., 2013), supporting our gene signatures. Among the differentially expressed genes we found the chloride/potassium symporter *Kcc2*, which is pivotal for the GABA switch from excitation to inhibition during neuronal maturation (Ge et al., 2006),_ selectively expressed in mature neurons as previously shown (Ge et al., 2006)._Interestingly, immature neurons in DG express genes for both GABA production (one of two GABA synthetase genes, *Gad1*, as shown (Zhao et al., 2010) and transportation (*Gat1*, figure S20), despite maturing to be primarily glutamatergic neurons (Ming and Song, 2011).

Evidence from diverse mammalian systems suggests that adult neurogliogenesis occurs in multiple non-canonical regions of the adult CNS (Feliciano et al., 2015). However, traditional methods, such as FISH, are limited in their ability to identify and fully characterize rare newborn cells. In particular, contradicting FISH evidence of a few marker genes suggests that progenitors in the adult spinal cord (SC) give rise either to only glia cells (Horner et al., 2000) or to both glia cells and neurons (Shechter et al., 2007). To systematically investigate neurogliogenesis in the adult SC, we applied Div-Seq and found a clear signature for dividing cells 7 days after EdU labeling (figure S19). Clustering analysis revealed a diverse population of newborn cells, in which the majority (54%) represented oligodendrocyte precursor cells (OPCs) (expressing *Sox10*), and the second largest population (29%) represented immature neurons (expressing *Sox11*) (**Figure 5A-B** and figure S19). Notebly, In the non-EdU labeled population we found mainly mature neurons (70%) and glia (30%) with only 4% OPCs and no immature neurons, demonstrating the need for Div-Seq to capture these rare cell types. All newborn neurons we detected expressed the GABA processing genes *Gad1* and *Gad2*, suggesting that newborn neurons in the SC are GABAergic (supporting previous observations (Shechter et al., 2007), **Figure 5B**).

**Figure 5.**
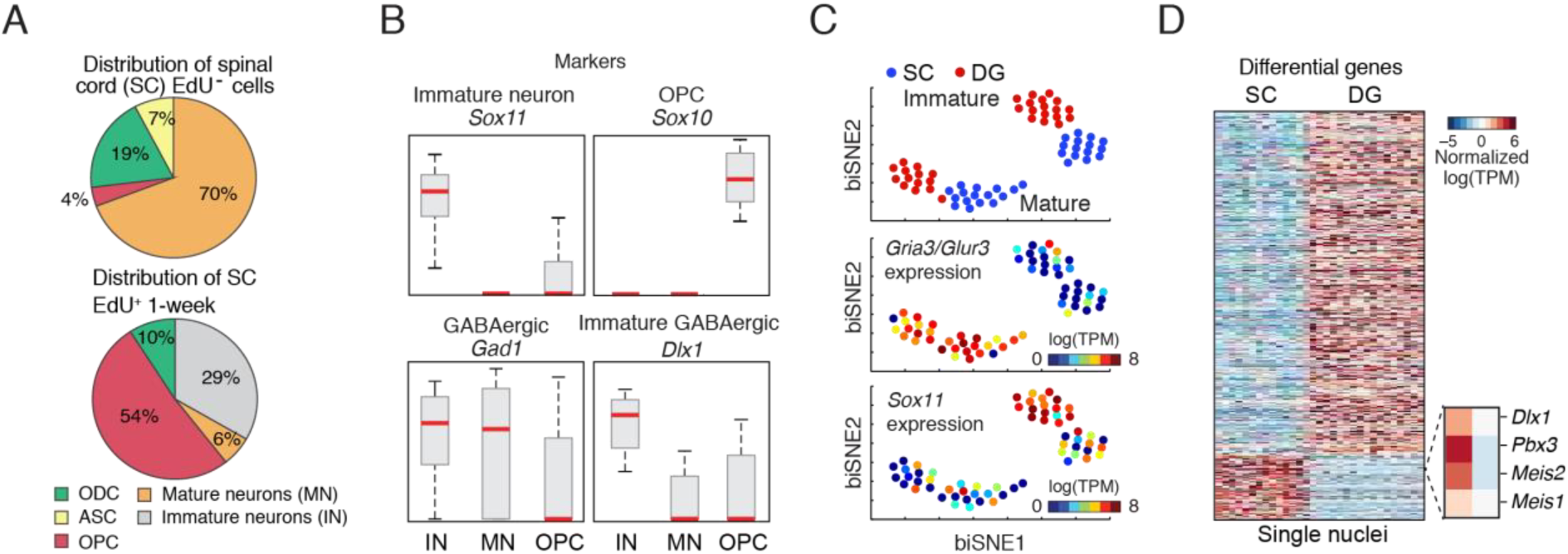
Adult newborn GABAergic neurons in the spinal cord revealed by Div-Seq. (A) Div-Seq in the spinal cord (SC) captures oligodendrocyte precursor cells and immature neurons. Shown is the distribution of cell types in the SC in non-EdU-labeled (top) and 7 days EdU labeled nuclei (bottom), assigned by clustering and marker gene expression. Oligodendrocyte precursor cells, OPC; Astrocytes, ASC; Oligodendrocytes, ODC. (B) Div-Seq captured immature neurons expressing marker genes of immature neurons (*Sox11*), GABAergic (*Gad1, Dlx1*) or OPCs (*Sox10*) marker genes. Box plots for immature neurons (IN), mature neurons (MN) and OPCs, shown as in **Figure 3G**. (C) Cells cluster primarily by maturation state and secondarily by region. All panels show biSNE 2D embedding of immature and mature neurons from both the SC and the DG. Nuclei are colored by tissue (top), by *Gria3/Glur3* mature marker expression levels (middle), or by *Sox11* (bottom). (D) Region specific gene expression. Heatmap shows the expression of genes specific to immature neurons in the SC (left) and DG (right), across single nuclei (t-test FDR<0.05, log-ratio>1, across all pairwise comparisons).

Comparison of immature and mature neurons in both the SC and the DG revealed that cells cluster primarily by maturation state and secondarily by region (**Figure 5C**), demonstrating genetic similarities between immature neurons independent of their origin within the CNS. However, focusing on immature neurons we identified differentially expressed genes (**Figure 5D**) specific to the DG (e.g. *Prox1*) and the SC (e.g. *Rex2*), respectively. In particular, we found three transcription factors, *Pbx3, Meis2*, and *Dlx1* co-expressed specifically in the SC but not in DG immature neurons. Previous reports showed that *Pbx, Meis*, and *Dlx* super-family factors interact (Rottkamp et al., 2008) and promote adult neurogenesis in the subventricular zone/olfactory bulb and dopaminergic fate specification (Petryniak et al., 2007); our data suggest that these factors may also play a regulatory role in adult neurogenesis in the SC. Taken together, the comparison of RNA signatures of newborn neurons in the SC and DG suggests that common molecular pathways cooperate with cell type-specific fate specifying factors to mediate adult neurogenesis across different brain regions.

In summary, we have shown how Nuc-Seq and Div-Seq open new avenues in the study of neuronal diversity and rare dynamic processes in the adult CNS. Nuc-Seq overcomes the harsh dissociation needed for single cell RNA-Seq, yet retains rich information required to make fine distinctions between cell types and states. Combined with intra-nuclear tagging, our nuclei profiling method enables the study of rare cell populations, as done in Div-Seq to capture proliferating cells. Future technology development may work to increase the sensitivity of Nuc/Div-Seq and to further extend the technologies by integrating with other techniques. For example, integration with droplet-based microfluidics may help to increase throughput, and the use of alternative labeling approaches such as immunostaining of transcription factors (Thomsen et al., 2016) or a recently published fluorescent “flash” tagging of dividing cells (Ludovic Telley) may broaden the range of cell types possible for investigation. Div-Seq’s ability to clearly identify and characterize rare cells in the spinal cord shows its significantly improved sensitivity compared to traditional methods. Nuc-Seq and Div-Seq can be readily applied to diverse biological systems, and may be especially helpful for studying transcriptional dynamics, the aging brain, fixed and frozen tissue, and time-sensitive samples such as human biopsies. Overall, our methods will help overcome broad challenges not only in neuroscience, but in many other biological systems as well.

## Acknowledgments

We thank R. Macrae, J.N. Campbell, A. Shalek, R. Satija, C. Dulak, S. Kadosch, N. Friedman, D. Gennert, O. Rosen, Z. Wang and the Zhang and Regev laboratories for support and discussions. N.H. is supported by HHMI and HHWF. M.H. is supported by the Human Frontier Science Program. This work was supported by the KCO, NIMH grant U01MH105960 (to F.Z. and A.R.), NIDDK grant 5R01DK097768-03 (to F.Z.). F.Z. is a NYSCF Investigator and A.R. is an HHMI Investigator. The authors have no conflicting financial interests.

